# Eukaryotic initiation factor 2α kinases regulate virulence functions, stage conversion, and the stress response in *Entamoeba invadens*

**DOI:** 10.1101/2022.03.03.482935

**Authors:** Heather A. Walters, Brenda H. Welter, Ronny Orobio-Hurtado, Harrison C. Moss, Martha A. Villano, William J. Sullivan, Lesly A. Temesvari

## Abstract

*Entamoeba histolytica* is a protozoan parasite that causes amoebic dysentery and liver abscess. This pathogen possesses a two-stage life cycle consisting of an environmentally stable cyst and a pathogenic amoeboid trophozoite. Since infection is acquired by ingestion of cysts from contaminated food and water, this parasite is prevalent in underdeveloped countries. A reptilian pathogen, *Entamoeba invadens*, which can encyst in culture, has long-served as a surrogate to study stage conversion. In the host, the *Entamoebae* must manage stress including nutrient deprivation and host immune pressure. In many systems, the stress response is characterized by down-regulation of translation, which is initiated by the phosphorylation of eukaryotic initiation factor-2 alpha (eIF2α). In mammalian cells, this phosphorylation is carried out by a family of eIF2α kinases. A canonical eIF2α translational control system exists in the *Entamoebae*; however, no eIF2α kinases have been characterized. In this study, we identified two eIF2α kinases in *E. invadens*, EiIF2K-A and EiIF2K-B. Their identity as eIF2α kinases was validated using a heterologous yeast system. We used an RNAi Trigger-mediated silencing system to reduce expression of EiIF2K-A, which also reduced expression of EiIF2K-B. Parasites with decreased kinase expression exhibited decreased phosphorylation of eIF2α and increased sensitivity to oxidative stress. Diminished kinase expression also correlates with an increased rate of encystation, a decreased the rate of excystation, and an increase in several virulence functions, erythrophagocytosis and adhesion to host cells. Taken together, these data suggest that EiIF2K-A and EiIF2K-B are authentic eIF2α kinases that may regulate the *Entamoeba* stress response.

**Importance:** *Entamoeba histolytica* is a human pathogen that causes dysentery and affects millions of people worldwide. This parasite possesses a two-stage life cycle: an environmentally stable cyst and the pathogenic trophozoite. Cysts are ingested from contaminated food and water; thus, this parasite in prevalent in underdeveloped countries. Current therapies commonly cause adverse side effects; therefore, new treatments are needed. In the host, *Entamoeba* experiences stress brought on, in part, by the host immune system. Understanding stage conversion and the stress response of this pathogen may lead to new drug therapies. Using the model organism, *E. invadens*, we identified two kinases, similar to those involved in stress and stage conversion in other systems. We determined that these kinases may regulate the oxidative stress response, stage conversion, and virulence. This work is significant as it will inform future studies on the life cycle and pathogenicity of the *Entamoeba* species.

## Introduction

*Entamoeba histolytica* is a human pathogen that causes amoebiasis and amoebic liver abscess, affecting millions of people worldwide and causing an estimated 55,000 deaths annually [1]. *E. histolytica* has a two-stage life cycle: the infectious cyst and the pathogenic amoeboid trophozoite. Latent cysts are ingested from fecally-contaminated food or water; thus, this parasite is prevalent in underdeveloped countries, where infrastructure, especially sanitation, is substandard. In 2015, 663 million people lacked access to clean drinking water and almost 1 billion people still practiced open defecation [2]. Additionally, amoebiasis is the leading cause of diarrheal disease in travelers returning to the US [1]. Considered together, these characteristics demonstrate that *E. histolytica* constitutes a significant global health problem.

After ingestion, cysts traverse the stomach and enter small intestine, where unknown cues trigger the excystation of trophozoites. These amoebae travel to the colon where infection can progress along several non-mutually exclusive routes. The trophozoites may establish a noninvasive infection, feeding on gut flora or host cells by phagocytosis. The parasites may also adhere to and degrade the gut epithelial lining, causing a diarrheal disease known as amoebic dysentery. Occasionally, the parasites breach the intestinal wall, enter the bloodstream, and establish extraintestinal infection in the liver (amoebic liver abscess), or more rarely, in the lungs and brain. In the large intestine, unknown signals trigger the conversion of trophozoites into environmentally stable mature cysts that are shed into the environment to facilitate host-to-host spread [3]. While navigating the human host, *E. histolytica* faces numerous stresses, such as nutrient deprivation, oxidative stress, nitrosative stress, and heat shock [4,5]. To survive, the parasite must surmount these damaging conditions.

*E. invadens*, a reptilian parasite, has served as a model to study stage conversion in this genus, because it readily encysts and excysts in culture [6,7,8]. Stage conversion is thought to be a response to stress encountered in the colon, and many of the features of the stress response overlap with those of stage conversion. For example, both heat shock proteins and cyst wall proteins are upregulated during heat shock in *E. invadens* [9]. Additionally, a eukaryotic type IIA topoisomerase II is upregulated during oxidative stress, heat shock, and encystation [10]. Given the importance of stress management during the parasite’s life cycle, stress response pathways may represent a novel targetable vulnerability. Thus, it is crucial to understand the molecular mechanisms that regulate the parasite stress response. Such information would provide significant insight into *Entamoeba* pathogenicity and would inform future studies focused on anti-parasitic drug design.

In most organisms, one branch of the stress response is characterized by the phosphorylation of a conserved serine residue in the alpha subunit of eukaryotic initiation factor-2 (eIF2α). eIF2α is a component of a ternary complex with GTP and the initiator methionyl transfer RNA (Met-tRNAi). This ternary complex binds the 40S ribosomal subunit, delivering Met-tRNAi for translation initiation. Phosphorylation of eIF2α during stress inhibits this activity, causing a sharp decline in global protein synthesis and preferential translation of a subset of mRNAs that encode stress-related regulators. This process of translational control allows cells to conserve resources and reconfigure gene expression to effectively counter stress. In mammalian cells, phosphorylation of eIF2α is regulated by a family of four eIF2α kinases (GCN2, PKR, PERK, HRI) that are activated in a stress-specific manner. GCN2 is activated by nutrient starvation, PKR is activated in response to viral infections, PERK is activated by misfolded proteins, and HRI is activated by heme starvation [11]. Although translational control, via eIF2α phosphorylation, was shown to exists in *E. histolytica* [4,5], no eIF2α kinases have been characterized in the *Entamoebae.*

In this study, we identified two putative eIF2α kinases, EiIF2K-A and EiIF2K-B in *E. invadens*. We used a heterologous yeast system [12,13] to confirm that EiIF2K-A and EiIF2K-B are *bona fide* eIF2α kinases. We used a Trigger-mediated silencing approach [14] to knock down expression of EiIF2K-A, which simultaneously reduced expression of EiIF2K-B. Parasites with reduced expression of these kinases exhibited decreased levels of phosphorylated eIF2α, a diminished ability to surmount oxidative stress, and altered rates of stage conversion. Furthermore, decreased kinase expression was correlated with an increase of two virulence functions, erythrophagocytosis and adhesion. Taken together, these data show that EiIF2K-A and EiIF2K-B are authentic eIF2α kinases that may be involved in the parasite stress response, stage conversion, and virulence.

## Results

### The *E. invadens* and *E. histolytica* genomes each encode two putative eIF2α kinases

Using the amino acid sequences of the four human eIF2α kinases, we searched the *E. invadens* genome (Amoebadb.org) for candidate sequences that contained hallmarks of eIF2α kinases [15]. We found two presumptive *E. invadens* eIF2α kinases, which we named EiIF2K-A (EIN_052050, formerly labeled EIN_033330) and EiIF2K-B (EIN_096010, formerly labeled EIN_059080), which share ~33.5% identity and ~48.1% similarity with each other within their kinase domains (Table S1). According to RNA sequencing data, reported as transcript abundance in transcripts per million (TPM) (Amoebadb.org), these kinases exhibit stage-specific expression. EiIF2K-A is predominantly expressed in trophozoites and at 48 h into encystation, while EiIF2K-B is only expressed during stage conversion, at low levels during encystation, and at higher levels during excystation [8]. Like the genome of *E. invadens*, the genome of the human pathogen, *E. histolytica*, also possesses two putative eIF2α kinases, which we named EhIF2K-A (EHI_035950) and EhIF2K-B (EHI_109700). The kinase domain of EiIF2K-A shares ~49.9% identity (~64.6% similarity) and ~28.1% identity (~43.9% similarity) with the kinase domains of EhIF2K-A and EhIF2K-B, respectively (Table S1). The kinase domain of EiIF2K-B shares ~31.1% identity (~45% similarity) and ~48.4% identity (~64.9% similarity) with the kinase domains EhIF2K-A and EhIF2K-B, respectively (Table S1). The kinase domains of EiIF2K-A and EiIF2K-B also share at least 16.57% identity and at least 28.57% similarity with the human eIF2α kinases (Table S1).

We aligned the putative *Entamoeba* kinases with other known eIF2α kinases, as well as with a control kinase, human CDK1, which does not belong to the family of eIF2α kinases (Fig 1). The *Entamoeba* kinases possess all eleven subdomains characteristic of the eIF2α kinase family, with highly conserved residues making them more closely related to eIF2α kinases than to the control kinase, CDK1 (Fig 1). The *Entamoeba* kinases share little homology with other members of this kinase family beyond the kinase domains (Fig 1). A phylogenetic analysis of eIF2α kinases showed that both EiIF2K-A and EiIF2K-B were more closely related to each other and to their *E. histolytica* counterparts (EhIF2K-A and EhIF2K-B) than to any of the other kinases. Additionally, the *Entamoeba* kinases were more closely related to PKR- and PERK-related kinases than to GCN2- or HRI-related kinases (Fig 2).

**Figure 1:**
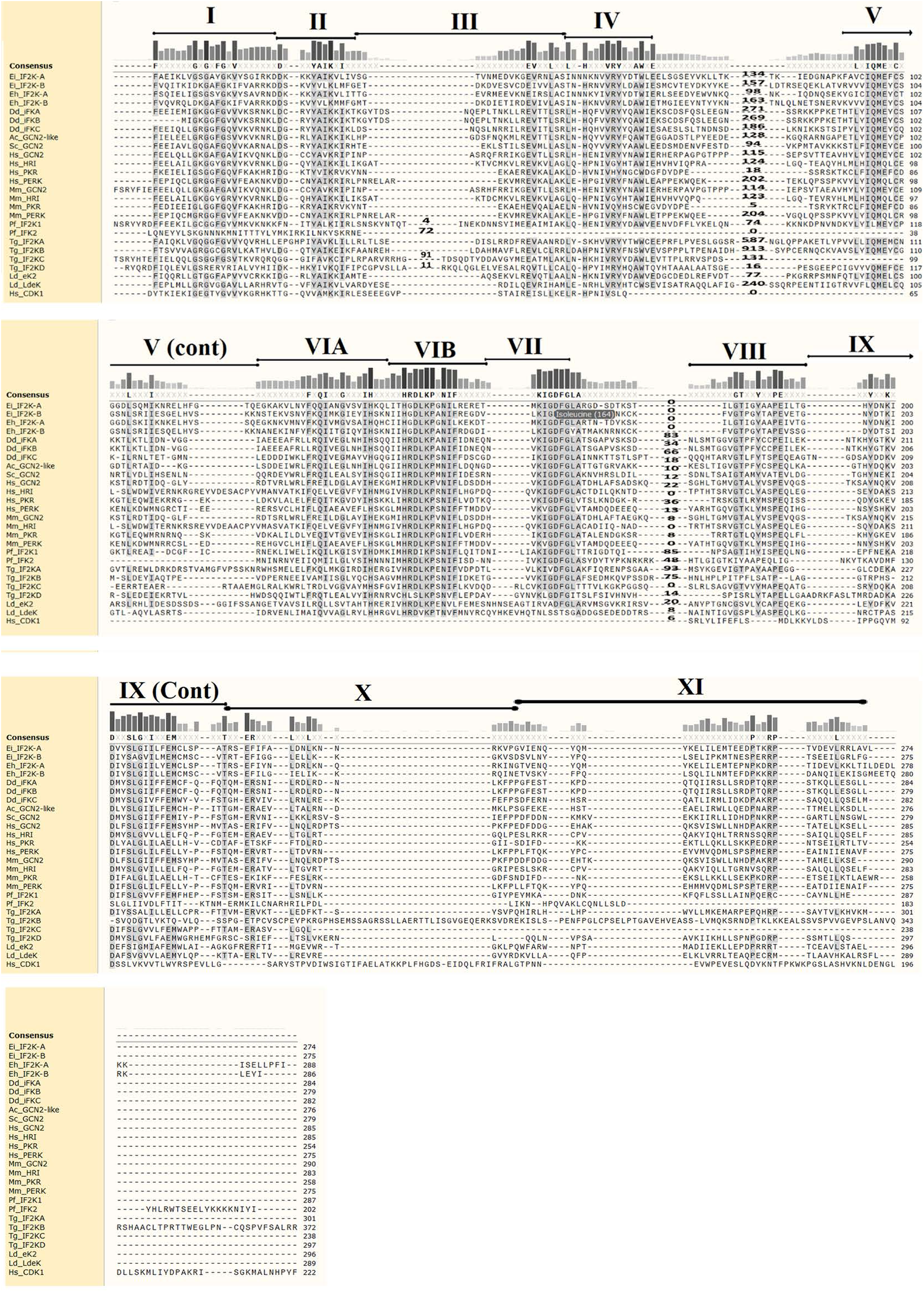
Alignment of kinase domains. Clustal W alignment of the catalytic domains of different eIF2α kinases. Roman numerals indicate each of the subdomains characteristic of a kinase. The grey scale bars represent level of residue conservation. Dashes represent gaps in the sequences that were used to maximize the alignment. The inserts between subdomains IV and V, and VII and VII were removed for clarity. The numbers between the dashes indicate the size of each insert removed. NCBI Accession numbers: *Entamoeba invadens*; EiIF2K-A (XP_004259781.1), EiIF2K-B (XP_004254115.1). *Entamoeba histolytica*; EhIF2K-A (XP_648932.2), EhIF2K-B (XP_652189.2). *Dictyiostelium discoideum*; Dd iFKA (Q558U1.1), Dd_iFKB (Q550L8), Dd_iFKC (Q75JN1). *Acanthamoeba castellani*; Ac_GCN2-like (L8HJ53). *Saccharomyces cerevisiae*; Sc_GCN2 (P15442). Human; Hs_GCN2 (Q9P2K8.3), Hs_HRI (Q9BQI3), Hs_PKR (P19525), Hs_PERK (Q9NZJ5), Hs_CDK1 (NP_203698). Mouse; Mm_GCN2 (NP_001171277.1), Mm_HRI (Q9Z2R9), Mm_PKR (Q03963), Mm_PERK (Q9Z2B5). *Plasmodium falciparum*; Pf_IF2K1 (XP_001348597.1), Pf_IFK2 (Q8I265). *Toxoplasma gondii*; Tg_IF2KA (S8F350), TgIF2K-B(ACA62938), Tg_IF2KC (AHM92904), Tg_IF2KD (AED01979.1). *Leishmania donovani*; Ld_eK2 (A0A0F7CYG9), Ld_LdeK (A9YF35)

**Figure 2.**
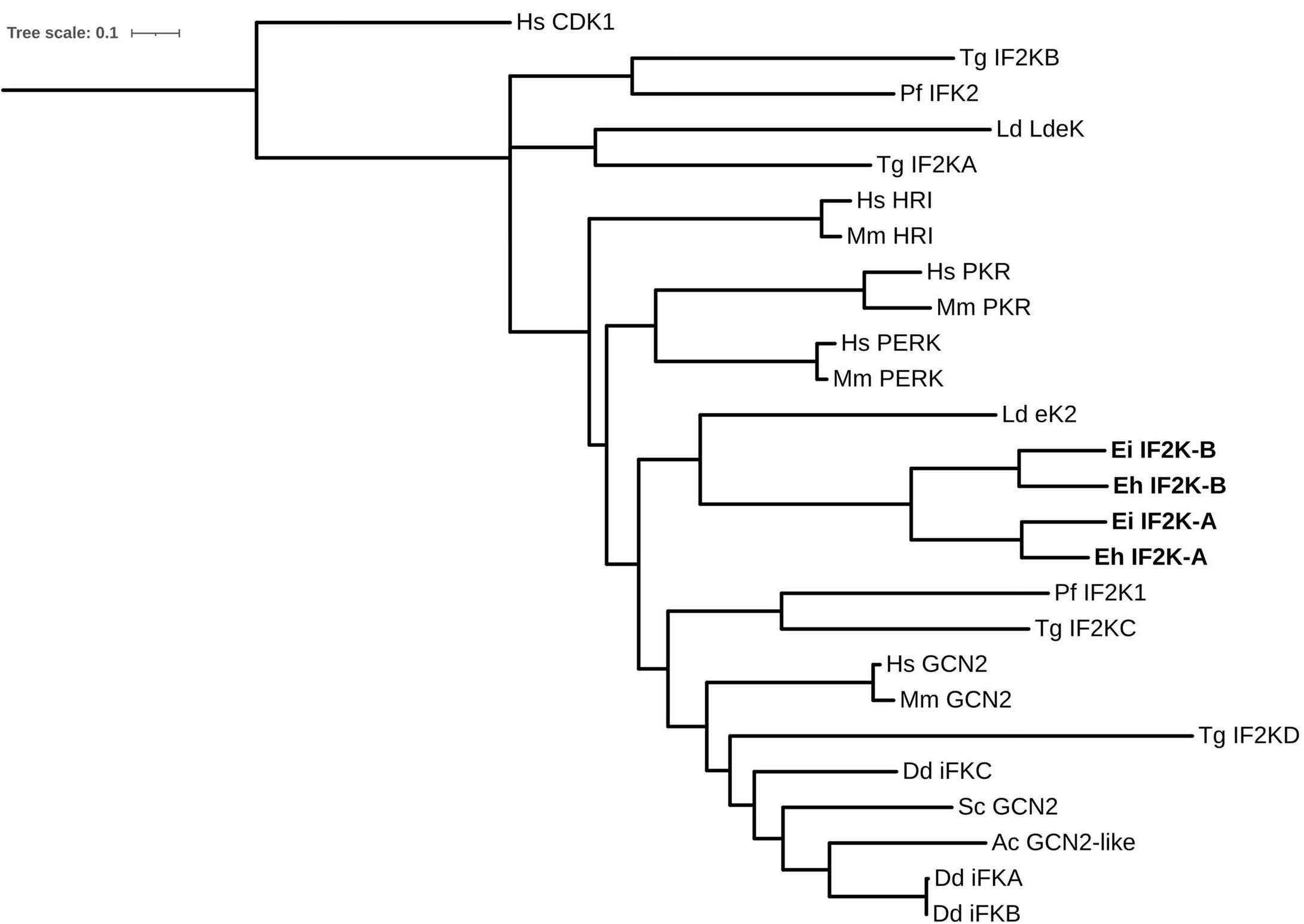
Phylogenetic tree of eIF2a kinases. The alignment of the catalytic domains of known eIF2a kinases and putative *Entamoebae* kinases was used to propose phylogenetic relationships by generating a tree. NCBI Accession numbers: *Entamoebae*; EiIF2K-A (XP_004259781.1), EiIF2K-B (XP_004254115.1), EhIF2K-A (XP_648932.2), EhIF2K-B (XP_652189.2). *Dictiostelium discoideum*; Dd iFKA (Q558U1.1), Dd iFKB (Q550L8), Dd iFKC (Q75JN1). *Acanthamoeba castellani*; Ac GCN2-like (L8HJ53). *Saccharomyces cerevisiae*; Sc GCN2 (P15442). Human; Hs GCN2 (Q9P2K8.3), Hs HRI (Q9BQI3), Hs PKR (P19525), Hs PERK (Q9NZJ5), Hs CDK1 (NP_203698). Mouse; Mm GCN2 (NP_001171277.1), Mm HRI (Q9Z2R9), Mm PKR (Q03963), Mm_PERK (Q9Z2B5). *Plasmodium falciparum*; Pf IF2K1 (XP_001348597.1), Pf IFK2 (Q8I265). *Toxoplasma gondii*; Tg IF2KA (S8F350), Tg IF2K-B (ACA62938) Tg IF2KC (AHM92904), Tg IF2KD (AED01979.1). *Leishmania donovani*; Ld eK2 (A0A0F7CYG9), Ld LdeK (A9YF35).

### EiIF2K-A and EiIF2K-B regulate phosphorylation of eIF2α in a heterologous system

To validate that EiIF2K-A and EiIF2K-B are eIF2α kinases, we utilized a yeast model system that uses the *Saccharomyces cerevisiae* strain, H1894, in which the sole endogenous eIF2α kinase is deleted. Exogenous expression of authentic eIF2α kinases in this yeast strain results in phosphorylation of endogenous eIF2α. [12,13,16]. A truncated cDNA encoding the catalytic domain of EiIF2K-A or EiIF2K-B, was inserted into the yeast expression vector, pYES-NT/C, which confers uracil prototrophy and allows for galactose-inducible expression of exogenous genes [13]. The pYES-NT/C plasmid also adds a polyhistidine tag to the N-terminus of the expressed protein. A pYES-NT/C vector lacking an insert (empty pYES) was used as a control plasmid and pYES2 plasmid harboring the active kinase domain of human PKR was used as a positive control [13]. A standard transformation protocol [17] was used to introduce the expression vectors into the H1894 strain and transformants were selected by growth on media that lacked uracil.

Expression of exogenous protein was induced by plating yeast cells on galactose-containing medium. Western blot analysis using an α-polyhistidine antibody demonstrated successful induction of exogenous protein expression with little to no expression prior to exposure to galactose (Fig 3A). We used western blot to assess the level of phosphorylated and total eIF2α in the transgenic yeast strains expressing EiIF2K-A, EiIF2K-B, empty pYES, or human PKR. Both *E. invadens* kinases phosphorylated endogenous yeast eIF2α (Fig 3B) demonstrating that EiIF2K-A and EiIF2K-B have eIF2α kinase activity.

**Figure 3:**
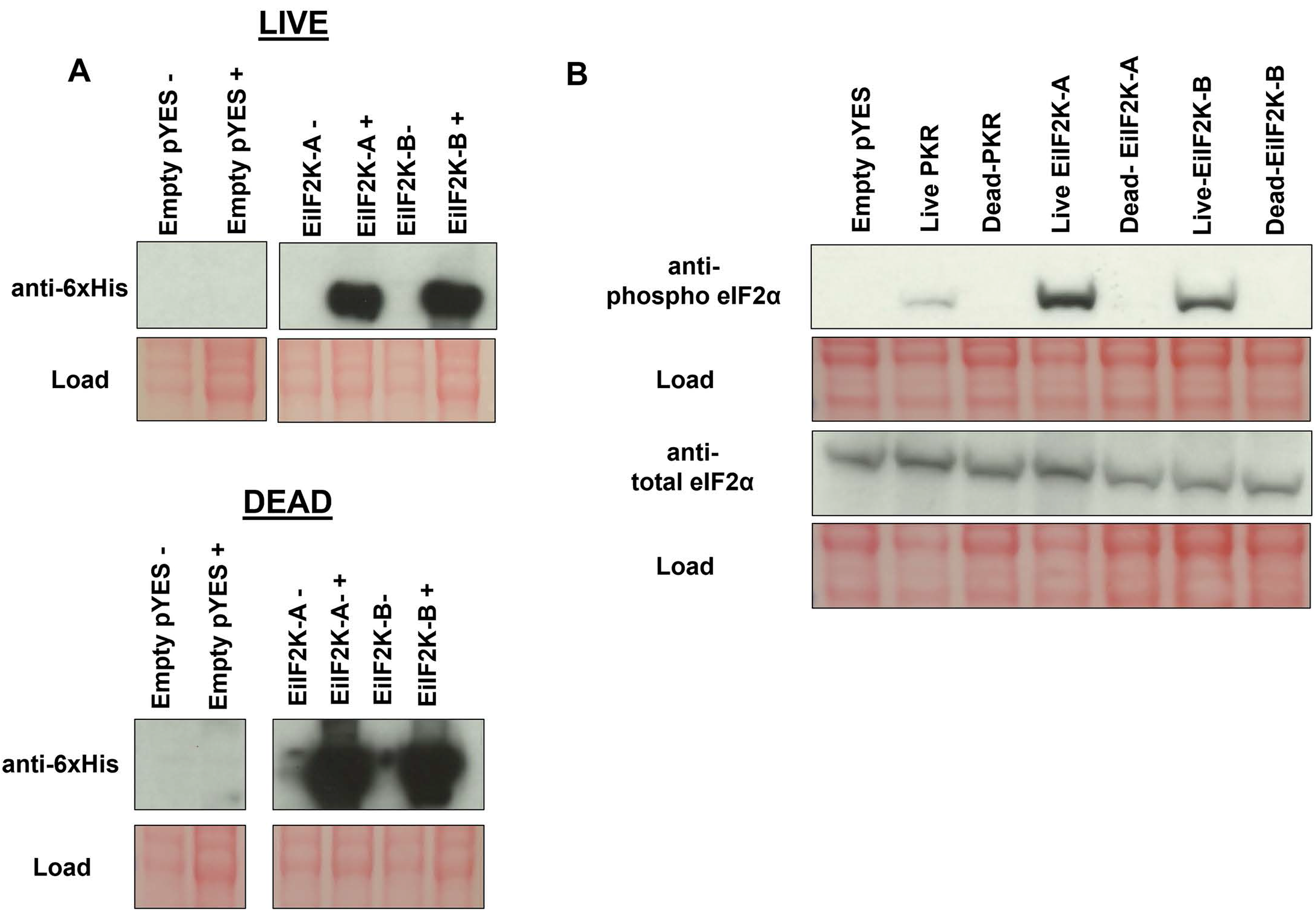
Expression of EiIF2K-A and EiIF2K-B kinase domains in a heterologous yeast system demonstrates kinase activity. Yeast strain H1894, which contains a genetic deletion of its sole eIF2α kinase, was transformed with the galactose-inducible pYES expression vector (empty pYES) or the same vector harboring wildtype (live) or mutated (dead) coding sequences for EiIF2K-A or EiIF2K-B kinase domains. For the dead kinases, a conserved lysine in each kinase subdomain II was mutated to arginine to create an inactive kinase. (A) Western blot, using anti-polyhistidine tag antibody, confirmed galactose-inducible protein expression. “-“ symbol represents strains grown on glucose as a carbon source, while “+” symbol represents strains grown on galactose as a carbon source. Expression of the EiIF2K-A or EiIF2K-B kinase domains (live or dead) was only evident when cells were grown on galactose. (B) Western blot showing the level of phosphorylated eIF2α and total eIF2α in H1894 yeast strain harboring empty pYES, a control eIF2α kinase, human PKR (pYES-PKR (live or dead) [13]), pYES-EiIF2K-A (live or dead), or pYES-EiIF2K-B (live or dead). Hyperphosphorylation of eIF2α was only observed in the yeast expressing live kinase domains. Ponceau red staining of membranes (red) indicate load.

EiIF2K-A and EiIF2K-B possess the conserved lysine in kinase subdomain II (Fig 1) that is critical for catalytic activity [18]. As a control, we mutated this key lysine residue to arginine and expressed these “dead” kinase domains in the H1894 yeast strain. Western blotting confirmed that dead kinases were incapable of phosphorylating yeast eIF2α (Fig 3B). These findings further support the notion that EiIF2K-A and EiIF2K-B are authentic eIF2α kinases.

### EiIF2K-KD parasites exhibit lower phospho-eIF2α levels and altered growth in nutrient-rich media than control parasites

We used a RNAi Trigger-mediated gene silencing approach to reduce the expression of EiIF2K-A [14]. Since the coding sequence of EiIF2K-A is large (2727 nucleotides), we subcloned a partial cDNA encoding amino acids 1-267, which contains the kinase domain, into the gene silencing Trigger plasmid [14]. Complete cDNAs are not required for efficient knockdown using this system [14]. Stable transfectants (EiIF2K-KD) were selected for and maintained by growth in the presence of neomycin. Parasites harboring the Trigger plasmid, with an insert encoding luciferase, an irrelevant protein, was used as a control (Trig Luc). Using this approach, we obtained substantial knockdown of EiIF2K-A mRNA levels in trophozoites as assessed by RT-PCR analysis (Fig S1A). Additionally, we measured EiIF2K-A expression during stage conversion in control and knockdown parasites. We found that EiIF2K-A mRNA was undetectable in control and knockdown parasites at 24 h (Fig S1B) and 48 h (Fig S1C) into encystation. EiIF2K-A mRNA was expressed at low levels in the control Trig Luc parasites at 2 h into excystation but was undetectable in the EiIF2K-KD parasites (Fig S1D). Since closely related genes may simultaneously be silenced by RNAi approaches [14], it was necessary to also measure expression of EiIF2K-B during growth and stage conversion. EiIF2K-A and EiIF2K-B share 32.64% identity within the kinase domain (Table S1). Consistent with published transcriptomic data [8], expression of EiIF2K-B was undetectable in control or mutant trophozoites growing in nutrient-rich medium (Fig S1A). On the other hand, the level of EiIF2K-B mRNA was reduced during both encystation (Fig S1B) and excystation (Fig S1D) in the EiIF2K-KD parasites compared to the Trig Luc control parasites, suggesting that reducing expression of EiIF2K-A simultaneously reduced expression of EiIF2K-B during stage conversion.

Previously, we showed that *E. histolytica* parasites possess a basal level of phosphorylated eIF2α, which increases after exposure to a subset of stressful conditions [4,5] and during encystation [4]. Therefore, we measured the level of phosphorylated eIF2α relative to total eIF2α in both trophozoites and encysting control and EiIF2K-KD parasites (Fig 4). In agreement with previously published data [4], Trig Luc control parasites exhibited a basal level of phosphorylated eIF2α, which increased at 48 and 72 h into encystation (Fig 4A, B). In contrast, parasites with diminished kinase expression displayed decreased, albeit slightly variable, levels of phosphorylated eIF2α in trophozoites and in encysting parasites. The most dramatic decrease in phosphorylation of eIF2α was observed in the mutant at 48 h into the stage conversion program. While mRNA levels of EiIF2K-A and EiIF2K-B are undetectable by RT-PCR in our knockdown cell line (Fig S1), there must be some remaining level of kinase mRNA expression as we see some phosphorylation of eIF2α in EiIF2K-KD parasites (Fig 4A, B). Overall, these data demonstrate that EiIF2K-KD parasites have a reduced capacity for phosphorylating eIF2α, supporting the identity of EiIF2K-A and EiIF2K-B as eIF2α kinases.

**Figure 4:**
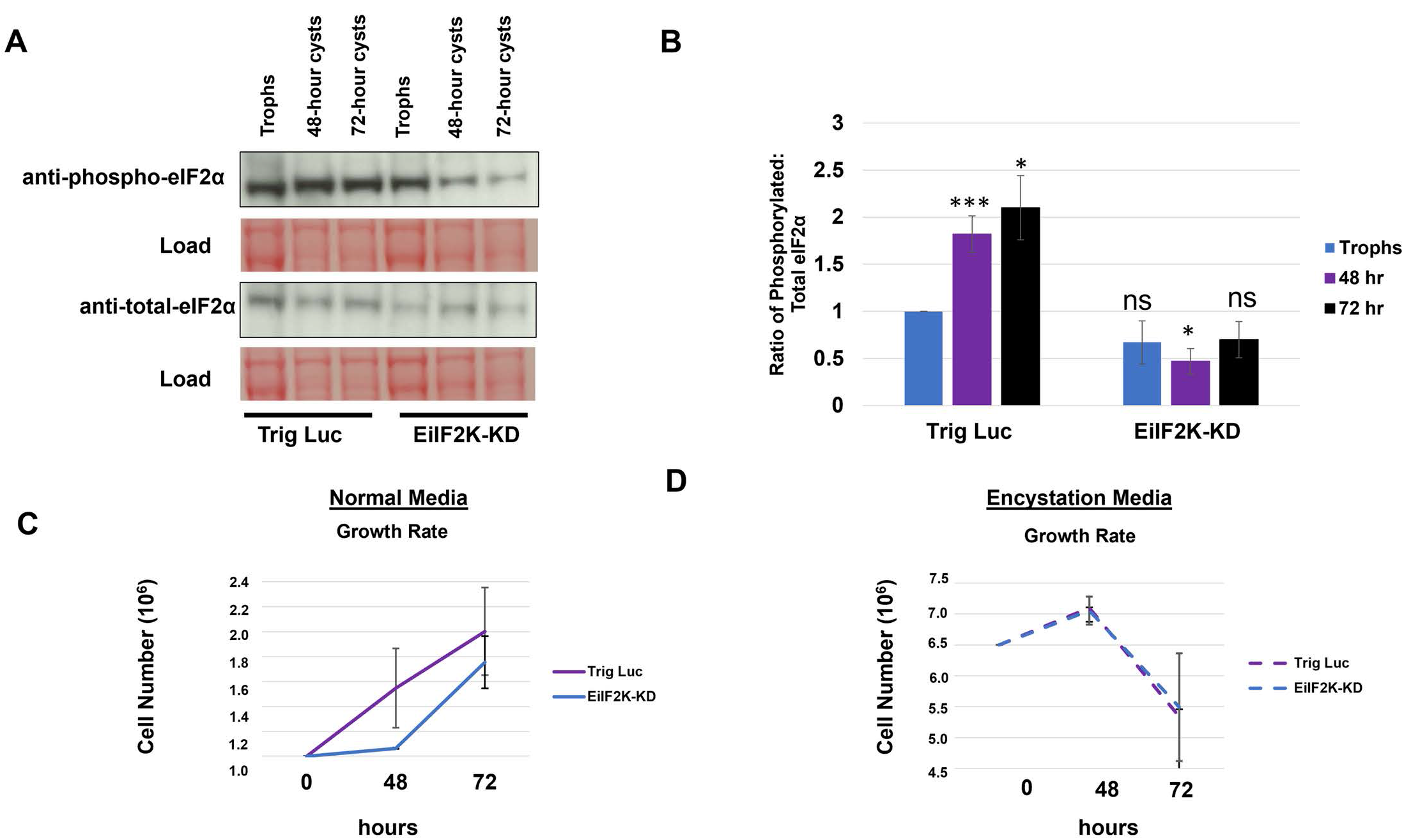
EiIF2K-KD parasites exhibit reduced phosphorylation of eIF2α and reduced growth in nutrient-rich medium. The level of phosphorylated and total eIF2α in Trig Luc and EiIF2K-KD parasites was measured during growth and encystation by western blotting using antibodies specific for phosphorylated or total or eIF2α. Levels of protein were quantified using scanning densitometry of bands on the same blot (Image J) and the ratio of phosphorylated eIF2α to total eIF2α was calculated after correcting for load. (A) Representative western blots for control (Trig Luc) or knockdown (EiIF2K-KD) trophozoites and encysting parasites. (B) Ratio of phosphorylated eIF2α to total eIF2α for Trig Luc or EiIF2K-KD trophozoites (blue), or 48-h cysts (purple), and 72-h cysts (black). The ratio for Trig Luc trophozoites was arbitrarily set to 1.0 and was used as the basis for comparison. During encystation, the ratio of phosphorylated eIF2α to total eIF2α increases in Trig Luc parasites (*P*<0.05). In all stages of the life cycle, the ratio of phosphorylated eIF2α to total eIF2α was generally decreased in EiIF2K-KD parasites compared to Trig Luc parasites. The most dramatic decrease in the ratio of phosphorylated:total eIF2α in was observed in the mutant at 48 h into the encystation program (*P*<0.05). (ns, not statistically significant). Data represent the mean ± standard error of at least 5 separate trials. Trophozoites were seeded into standard nutrient-rich culture media (1×10^6^ initial inoculum) (C) or into encystation media (6.5×10^6^ initial inoculum) (D) for 48 or 72 h. At each time point, parasites were enumerated using trypan blue exclusion and light microscopy. (C) EiIF2K-KD parasites exhibit an initial lag in growth when seeded into nutrient-rich medium, but eventually show an increased growth rate between 48 and 72 h compared to Trig Luc parasites. (D) EiIF2K-KD and Trig Luc parasites show no difference in growth rate at 48 or 72 h. Data represent the mean ± standard error of at least 3 separate trials.

We also measured the growth rate of Trig Luc and EiIF2K-KD parasites in both standard nutrient-rich medium and nutrient-poor/low osmolarity encystation medium for up to 72 h post-inoculation (Fig 4). EiIF2K-KD parasites exhibited a lag in growth when seeded into nutrient-rich medium, but eventually exhibited a higher rate of growth than control parasites (Fig 4C). This growth phenotype, exhibited by EiIF2K-KD parasites in nutrient-rich medium, was likely the result of reduced EiIF2K-A expression as it is the only kinase expressed in trophozoites. On the other hand, there was no difference in the growth kinetics of the mutant in encystation medium when compared to that of the control parasites (Fig 4D). The decrease in parasite number during incubation in encystation medium (Fig 4C, D) is typical as a fraction of the population loses viability instead of encysting.

### EiIF2K-KD parasites have altered rates of stage conversion

To elucidate the role of the kinases in stage conversion, we measured the rate of encystation and excystation in both Trig Luc and EiIF2K-KD parasites. Encystation was induced by inoculating parasites into nutrient-poor/low osmolarity encystation media. Hallmarks of encystation include the accumulation of a chitin-rich cell wall and a reduction in cell size [19]. To assess encystation, we used flow cytometry [19] and Congo Red staining to track the accrual of chitin as well as changes in cell size. In the EiIF2K-KD parasites, the percent of parasites that had encysted was significantly higher at 48 h post inoculation, but not at 72 h post inoculation, when compared to control parasites (Fig 5A). This suggests that the rate, but not the overall efficiency, of encystation is higher in parasites with diminished kinase expression. Since both kinases are expressed during encystation, we cannot discern if the encystation phenotype is due to loss of one or both kinases. To induce excystation, cysts were incubated in excystation media, which restores nutrients and osmolarity and contains bile salts to mimic passage through the host digestive system [8]. EiIF2K-KD parasites exhibited a significantly lower rate of excystation compared to control parasites at 2 h and 8 h into the excystation program (Fig 5B). Since EiIF2K-B is the only kinase expressed during excystation, we posit that reduced EiIF2K-B expression was responsible for this excystation phenotype.

**Figure 5:**
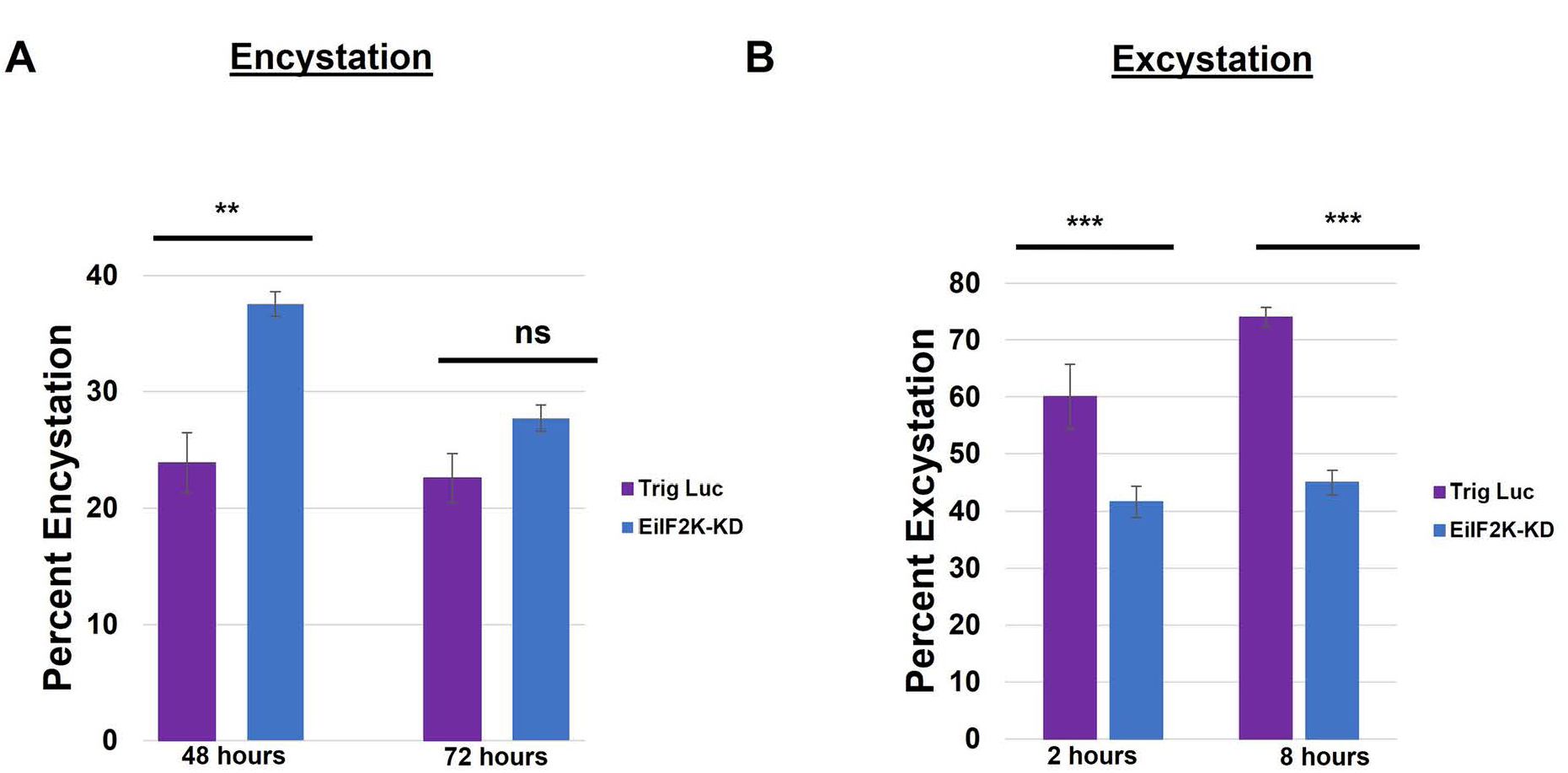
The rate of encystation is increased and the rate of excystation is decreased in EiIF2K-KD parasites. (A) Trig Luc and EiIF2K-KD parasites were induced to encyst for either 48 or 72 h. Mature cysts were stained with Congo Red and quantified using flow cytometry. A higher percent of EiIF2K-KD parasites encysted by 48 h and at 72 h compared to Trig Luc parasites. However, the increase was only statistically significant at 48 h (*P*<0.01), suggesting that the mutants have a higher initial rate of encystation but not a higher efficiency of encystation. (B) Trig Luc cells and EiIF2K-KD cysts were induced to excyst by incubation in excystation media for 2 or 8 h. The number of mature cysts was quantified before and after excystation and the decrease in the number of cysts represented the fraction (percent) of parasites that had excysted. The excystation rate of EiIF2K-KD was significantly (*P*<0.001) lower than that of Trig Luc parasites at both 2 and 8 h. Data represent the mean ± standard error of at least 3 separate trials.

### EiIF2K-KD trophozoites are more susceptible to oxidative stress

Previously, we demonstrated that *E. histolytica* phosphorylates eIF2α in response to several different stressful conditions including oxidative stress [4]. As such, EiIF2K-A, the sole kinase expressed in trophozoites, may be responsible for countering oxidative stress. Suresh et al., (2016) demonstrated that exposing *E. invadens* parasites to 4 mM H_2_O_2_ for 1 h induced oxidative stress, as evidenced by detachment and rounding of parasites, while maintaining ≥90% viability [14]. However, the level of phospho-eIF2α in H_2_O_2_-treated *E. invadens* trophozoites has not been examined. Therefore, we exposed wildtype (WT) *E. invadens* trophozoites to ddH_2_O (diluent) or 4 mM H_2_O_2_ for 1 h at 25°C and measured levels of total and phosphorylated eIF2α by western blotting (Fig S2A). Phosphorylation increased in parasites treated with 4 mM H_2_O_2_ compared to the unstressed control. To ascertain if EiIF2K-A regulates the response to oxidative stress in trophozoites, we measured the viability of WT, Trig Luc, and EiIF2K-KD parasites exposed to 4 mM H_2_O_2_; however, we observed no difference in viability (Fig S2B). Therefore, we used a higher concentration of H_2_O_2_ that could reduce viability of WT *E. invadens* parasites. WT, Trig Luc, and EiIF2K-KD parasites were exposed to 1 M H_2_O_2_ for 1 h at 37°C. This treatment caused approximately 30% parasite death in WT parasites, 40% parasite death in Trig Luc parasites (Fig 6A) and approximately 65% parasite death in EiIF2K-KD parasites. The statistically significant reduction in viability in EiIF2K-KD parasites supports the notion that EiIF2K-A may regulate the response to oxidative stress in *E. invadens* trophozoites.

**Figure 6:**
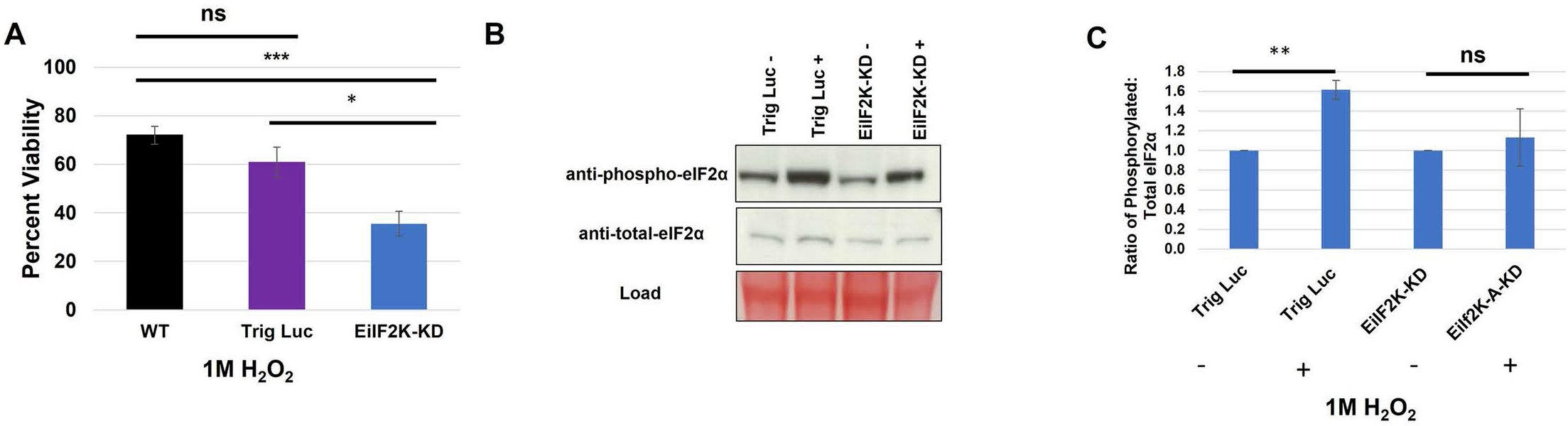
EiIF2K-KD trophozoites are more susceptible to oxidative stress. Wildtype (WT), Trig Luc, or EiIF2K-KD trophozoites were exposed to 1M H_2_O_2_ for 1 h at 25°C. (A) Viability was assessed using trypan blue exclusion and a Luna Automated Cell Counter. EiIF2K-KD parasites were significantly less viable when exposed to oxidative stress compared to WT and Trig Luc parasites (*P*<0.05). (B) Representative western blot showing the level of phosphorylated and total eIF2α in Trig Luc or EiIF2-KD cells before (-) and after (+) H_2_O_2_-treatment. (C) Western blotting was used to measure levels of total and phosphorylated eIF2α in parasites exposed to ddH_2_O or 1 M H_2_O_2_ for 1 h at 25·C. Levels of protein were quantified using scanning densitometry of bands on the same blot (Image J) and the ratio of phosphorylated eIF2α to total eIF2α was calculated after correcting for load. Trig Luc parasites exposed to 1 M H_2_O_2_ exhibited significantly higher (*P*<0.01) levels of phosphorylated eIF2α compared to controls, while EiIF2K-KD parasites exposed to the same conditions did not exhibit significantly increased levels of phosphorylated eIF2α. Data represent the mean ± standard error of at least 3 separate trials.

EiIF2K-KD parasites may be more susceptible to oxidative stress because of their reduced capacity to phosphorylate eIF2α. Thus, we used western blotting to measure the levels of phosphorylated and total eIF2α in control and EiIF2K-KD parasites exposed to H_2_O_2_. The ratio of phosphorylated eIF2α to total eIF2α increased significantly in stressed Trig Luc parasites, but not in EiIF2K-KD parasites (Fig 6B, C). There is a slight increase in phosphorylation of eIF2α in EiIF2K-KD parasites treated with 1M H_2_O_2_ compared to those parasites treated with ddH_2_O. Currently, there are no methods to knockout genes in the *Entamoebae*. Therefore, there remains some level of kinase expression in our EiIF2K-KD parasites, which may respond to stress. Overall, these data support the identity of EiIF2K-A as an authentic kinase and emphasize the importance of the eIF2α mechanism in parasite stress management.

### EiIF2K-KD trophozoites exhibit increased virulence functions

To discern the effect of decreased EiIF2K-A expression on parasite virulence, we measured two key virulence functions: erythrophagocytosis and adhesion to host cells. Trig Luc and EiIF2K-KD trophozoites were exposed to human red blood cells (hRBCs) for 10 min, after which uptake of heme was quantified spectrophotometrically [20]. Adhesion was measured by quantifying the degree to which fluorescently-labeled parasites could adhere to a fixed monolayer of Chinese hamster ovary (CHO) cells [21]. EiIF2K-KD parasites exhibited significantly increased phagocytosis (Fig 7A) and adhesion (Fig 7B), which suggests that EiIF2K-A may directly or indirectly modulates virulence functions, as EiIF2K-A is the only eIF2α kinase expressed by trophozoites

**Figure 7:**
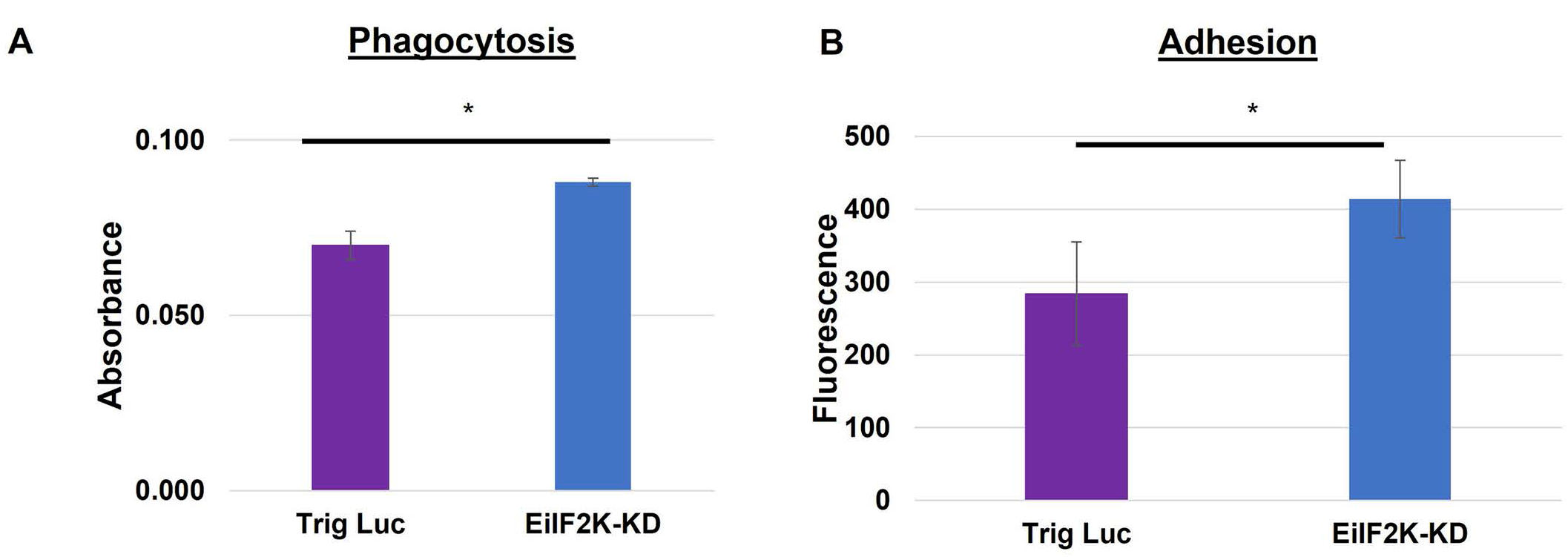
Erythrophagocytosis and adhesion are increased in EiIF2K-KD parasites. (A) Trig Luc or EiIF2K-KD parasites were incubated with human red blood cells (hRBCs: amoeba ratio; 100∶1) for 10 min, lysed, and spectrophotometrically analyzed for internalized heme at 405 nm. Amoebae with reduced expression of EiIF2K-A exhibited increased phagocytosis of hRBCs. The data represent the mean ± standard error. of at least 3 separate trials (*P*<0.05). (B) Calcein AM-stained control or mutant parasites were incubated with fixed monolayers of Chinese Hamster Ovary (CHO) cells for 30 min. Unadhered parasites were rinsed off the monolayer of CHO cells and the level of adhesion (calcein-AM fluorescence) was measured by spectrofluorimetry using an excitation wavelength of 485 nm and an emission wavelength of 528 nm. EiIF2K-KD trophozoites exhibited significantly higher adhesion to host cells than Trig Luc trophozoites (*P*<0.05). Data represent the mean ± standard error of at least 3 separate trials.

## Discussion

This is the first study to characterize eIF2α kinases in the *Entamoebae.* We used a heterologous yeast system to show that EiIF2K-A and EiIF2K-B are authentic eIF2α kinases. Using an established RNAi silencing approach [14], we knocked down both kinases using a single Trigger-EiIF2K-A plasmid. We found that EiIF2K-KD parasites were more susceptible to oxidative stress and exhibited increased virulence functions (erythrophagocytosis and parasite-host cell adhesion). We also observed an increased rate of encystation and decreased rate of excystation in EiIF2K-KD parasites. Due to the stage-specific expression patterns of these kinases, we posit that EiIF2K-A may regulate phenotypes observed in trophozoites, while the excystation phenotype may be due to the loss of EiIF2K-B. Overall, this study advances our knowledge about the stress response and stage conversion in *Entamoeba* species.

In mammalian cells, phosphorylation of eIF2α is regulated by a family of four eIF2α kinases that are activated in a stress-specific manner. The ability of the kinases to respond to various stresses rely on regulatory domains. Interestingly, in several protozoan parasites [13, 22, 23, 24], the eIF2α kinases possess divergent regulatory domains, suggesting that protozoan eIF2α kinases may respond differently to environmental stress than their mammalian counterparts [11]. Currently, it is not possible to predict, by sequence-analysis, the types of stresses to which the *Entamoeba* kinases will respond.

Nevertheless, we demonstrate that trophozoites with reduced EiIF2K-A expression are more susceptible to at least one stressful condition, oxidative stress. EiIF2K-KD parasites were less viable in the presence of high concentrations of H_2_O_2_ and possessed decreased levels of phosphorylated eIF2α when compared to control parasites (Fig 6A). This is not surprising since Hendrick et al. [4] demonstrated that oxidative stress induces the phosphorylation of eIF2α in *E. histolytica*. Likewise, Augusto et al. [25] knocked out an eIF2α kinase in *T. gondii*, TgIF2K-B, and found that null parasites had an impaired response to oxidative stress. To further illuminate the stress-specific response of this kinase, it will be necessary to assess the ability of the EiIF2K-KD cells to survive other stressful conditions. Additionally, examining the transcriptome and translatome of EiIF2K-KD and control parasites under oxidative stress would help determine if these eIF2α kinases directly regulate the stress response of *E. histolytica*.

EiIF2K-KD parasites exhibited no growth phenotype in encystation medium and a transient lag in growth in nutrient-rich medium (Fig 4C,D). This is unlike *Trypanosoma cruzi* parasites lacking the eIF2α kinase, TcK2, which exhibit a growth deficiency [26]. However, the *E. invadens* phenotype is similar that of *Leishmania donovani* parasites expressing a dominant negative version of a GCN2-like kinase, which do not exhibit a growth defect [22]. It is possible that in *L. donovani,* multiple eIF2α kinases share redundant functions and the loss of one kinase is compensated by other related kinases. However, in *E. invadens,* EiIF2K-A is the only eIF2α kinase expressed in trophozoites ([8] and the current study). Thus, compensation by related kinases may not be possible in the trophozoite stage of this parasite.

EiIF2K-A is expressed in trophozoites and decreases during initial encystation [8]. Since decreased EiIF2K-A expression correlates with initiation of encystation, it is conceivable that EiIF2K-KD parasites are primed to encyst. In support of this, the encystation rate of the EiIF2K-KD parasites was significantly higher than that of control parasites at 48 h (Fig 5A). If EiIF2K-KD parasites are primed to encyst, they may exhibit early expression of encystation-specific genes, which, in turn, could lead to an increased rate of encystation, but not necessarily an increased efficiency. To gain further insight into the relationship between eIF2α kinase expression and encystation, it will be necessary to define the cyst-specific translatome, perhaps by ribosome-profiling (Ribo-seq) [27], in EiIF2K-KD trophozoites.

The excystation rate of EiIF2K-KD parasites was significantly decreased (Fig 5B). EiIF2K-B is expressed at low levels during encystation, and at high levels during excystation [8]. Therefore, we hypothesize that the excystation phenotype may be due to the loss of EiIF2K-B, as it is the only eIF2α kinase expressed during excystation. At present, we cannot determine if the encystation phenotype is due to loss of EiIF2K-A, EiIF2K-B, or both. To understand the exact roles of EiIF2K-A and EiIF2K-B in stage conversion, it will be essential to knock down each gene individually and evaluate stage conversion.

Previously, we demonstrated that the level of phosphorylated eIF2α increases significantly during encystation [4]. Thus, it was not surprising that EiIF2K-A and/or EiIF2K-B may play a role in stage conversion in *E. invadens*. Likewise, eIF2α kinases play roles in stage conversion in other protozoa. For instance, phosphorylation of eIF2α increases during stage conversion or differentiation of *Trypanosoma cruzi* [26, 28], *T. gondii* [29], and *Plasmodium falciparum* [23]. It was unanticipated that reduced phosphorylation of eIF2α would correlate with an *increased* rate of encystation in *E. invadens*. Perhaps in the *Entamoebae*, precise timing of translation is necessary to control the rate of encystation in such a way as to guarantee the accurate conversion of trophozoites into environmentally-stable cysts. Without the kinases that control phosphorylation of eIF2α, the rate of translation becomes unbridled, and the rate of encystation becomes unregulated.

EiIF2K-KD parasites exhibited increased erythrophagocytosis and adhesion to host cells, which are two important virulence functions (Fig 7). These data suggest that EiIF2K-A, the only eIF2α kinase expressed in trophozoites [8], may directly or indirectly regulate erythrophagocytosis and adhesion. Similarly, *T. gondii* parasites lacking one eIF2α kinase, TgIF2K-B, were more virulent *in vivo* [25]. Given the role of eIF2α kinases in the management of translation, one explanation for increased parasite virulence functions is dysregulated translation of genes that control virulence.

It may be argued that an increase in virulence functions or the rate of encystation in the EiIF2K-KD parasites implies that this kinase is not a suitable target for anti-parasitic drug design. However, increased sensitivity to oxidative stress in the EiIF2K-KD parasites supports its potential as a drug target. The *Entamoebae* are microaerophilic. Therefore, to survive in the host, these parasites must preserve intracellular hypoxia within oxygenated host tissues, such as the liver, and surmount attacks on cellular homeostasis by reactive oxygen species originating from the host immune response [30]. Thus, it is conceivable that disabling EiIF2K-A would simultaneously restrict the ability of the pathogen to endure in the host. In support of this, genetic loss of the eIF2α kinases, PERK and GCN2, in immortalized mouse fibroblasts and human tumor cells increases their susceptibility to oxidative stress [31].

The eIF2α kinases are also implicated in human pathologies including cancer [32], diabetes [33], and neurodegenerative disorders [34] and are the subject of intense study because they represent logical targets for the design of therapies. For instance, overactivation of PERK has been associated with neurological disorders such as Parkinson’s Disease and Alzheimer’s Disease [34]. It has been found that the compound, LDN-0060609, significantly inhibits PERK-mediated phosphorylation of eIF2α in rat astrocytes, which suggests that it may be a suitable drug for the treatment of neurological diseases [35]. Targeting the eIF2α-based regulation of translation in protozoan parasites is also underway. For example, pharmacological inhibition of PK4 in *P. falciparum* with the PERK inhibitor, GSK2606414, blocks parasite differentiation and reduces artemisinin-induced latency [36]. Inhibition of PERK-like eIF2 kinase, TgIF2K-A, in *T. gondii*, with the same inhibitor, blocked multiple steps of the tachyzoite lytic cycle and lowered the rate of bradyzoite differentiation [37]. Finally, GSK2606414 reduced *Leishmania amazonensis* infection of macrophages [38]. Together, with the data presented in this study, these encouraging results in other pathogens support the potential for the *Entamoeba* eIF2α kinases to serve as targets for drug inhibition.

## METHODS

### Protein Alignment and phylogenetic analysis

The kinase domains sequences of the four putative *Entamoebae* kinases were obtained from UniProt [38]. Sequences were also analyzed using ScanProsite [39] to identify the kinase domains and to search for other possible domains and motifs. The catalytic domains of the four putative kinases were aligned with the kinase domains of previously characterized eIF2a kinases, using the Clustal W algorithm with the standard parameters with SnapGene (Version 5.2.1; GSL Biotech, LLC., San Diego, CA, USA). The software, Jalview [40], was used to remove the inserts with high length variability for clearer visualization. A phylogenetic analysis was performed using the unedited alignment and the website Méthodes et Algorithmes pour la Bio-informatique LIRMM (http://www.phylogeny.fr/index.cgi) [41]. The Newick format of the phylogeny was imported into the Interactive Tree of Life (iTOL) (itol.embl.de) [42] to generate the visual tree. All webpages and applications were used with the standard settings for each step. The tree was rooted to the more distantly related sequence of the pool (CDK1).

### Strains and Culture conditions

*Entamoeba invadens* (strain IP-1) was cultured axenically in TYI-S-33 medium in 15 mL glass screw cap tubes or 25 cm^2^ culture flasks at 25°C [44]. Parasites were passaged into fresh media every 7 days. Chinese hamster ovary (CHO) cells were cultured in DMEM supplemented with 10% FBS, PenStrep, and HEPES in 25 cm^2^ culture flasks at 37°C.

To generate a plasmid to reduce expression of EiIF2K-A, PCR was employed to amplify the kinase domain of EiIF2K-A using genomic DNA as a template and gene-specific primers (Table S2). The primers also added AvrII restriction sites to the 3’ and 5’ ends, which facilitated subcloning into the Trigger plasmid [14] (kind gift of Dr. Upinder Singh, Stanford University). Successful subcloning was confirmed by sequencing.

*E. invadens* was transfected by electroporation as described [43], with minor modifications. Briefly, two 25 cm^2^ flasks containing log-phase trophozoites were iced for 15 min to release adherent parasites. The parasites were collected by centrifugation at 500 × *g* for 5 min and washed with 20 mL ZM phosphate buffered saline (PBS) buffer (132 mM NaCl, 8 mM KCl, 8 mM NaPO_4_, 1.5 mM KH_2_PO_4_). Parasites were pelleted by centrifugation at 500 × *g* for 5 mins and resuspended in 1.6 mL complete ZM PBS buffer (ZM PBS with 0.5 mM Mg(CH_3_COO)_2_ • 4H_2_O and 0.09 mM CaCl_2_). Eight hundred µL of parasite suspension was combined with 150 µg plasmid DNA and electroporated in a 0.4 cm cuvette with two pulses at 1.2 kV and 25 µF using a BioRad Gene Pulser II. Parasites were transferred to 15 mL culture tubes containing 13 mL TYI-S-33 and allowed to recover for 48 h. Neomycin selection was added gradually at 5 µg/mL each week until a concentration of 50 μg/mL was reached.

To assess expression of EiIF2K-A and EiIF2K-B, RNA was extracted from trophozoites or cysts using TRIZOL (ThermoFisher; Waltham, MA). Two μg of total RNA was treated with RQ1 DNase enzyme (Promega; Madison, WI) per manufacturer’s instructions. Treated RNA was used to synthesize cDNA using the Invitrogen Superscript III First Stand Synthesis kit per the manufacturer’s instruction. (ThermoFisher). One µL of cDNA was used as template and PCR was carried out using EiIF2K-A specific primers or EiIF2K-B gene specific primers (Table S2). We confirmed that these primers do not cross-react to amplify both genes. EIN_327460 was used as an internal load control for analysis of gene expression in trophozoites and EIN_162500 was used as an internal load control for analysis of gene expression in cysts (see Table S2).

### Analysis of eIF2 kinase function in yeast

The coding sequence of the kinase domain of EiIF2K-A (1313 bp) and EiIF2K-B (1397 bp) was synthesized and ligated into the pYES-NT/C plasmid (ThermoFisher, kind gift of Dr. William Marcotte, Clemson University) by Genscript (Piscataway, NJ USA), using the restriction enzyme sites BamHI and NotI. The resulting construct contained the kinase domains in-frame with a N-terminal poly-histidine tag, which was confirmed by sequencing. To generate an inactive kinase, the conserved lysine in kinase subdomain II (EiIF2K-A, position 43 and EiIF2K-B, position 45), was mutated to arginine using the Phusion Site Directed mutagenesis kit (Thermo-Fisher) and mutagenic primers (Table S2). Successful mutagenesis was confirmed by sequencing. Live and dead human PKR kinase domains in yeast expression plasmid pYES2 (controls) were kind gifts from Dr. Ronald Wek (Indiana University School of Medicine).

The pYES-NT/C or pYES2 plasmids encoding the active or dead kinase domain, or no gene product (empty pYES-NT/C), were introduced into *Saccharyomyces cerevisiae* strain H1894 (*MATa ura3-52 leu2-3 leu2-112 gnc2∆ trp1 ∆-63*), which lacks the sole yeast eIF2α kinase, GNC2 [12,16]. Yeast was cultured at 30°C on YPAD agar plates containing 2% (w/v) glucose prior to transformation. Yeast transformation was carried out as described [17]. Since pYES-NT/C confers uracil prototrophy to transformants, selection was carried out by plating transformed yeast cells on agar plates containing synthetic dropout medium (SD) (Sigma-Aldrich St. Louis, MO, USA) (without uracil), and 2% (w/v) glucose, and growing them overnight at 30°C.

To induce expression of exogenous protein, cells from each transformed yeast strain were inoculated into liquid SD medium containing 2% (w/v) raffinose and 10% (w/v) galactose [13] and grown overnight at 30°Cprior to western blotting.

### Western blotting

Western blotting of whole cell lysates was used to assess the expression of kinases in yeast or the level of total and phosphorylated eIF2α in yeast or in *E. invadens*. For yeast, cell lysates were prepared as described [44]. Briefly, 1.89 × 10^7^ yeast cells were pelleted by centrifugation at 3,000 × *g* for 5 min. To prepare yeast for lysis, cells were resuspended in 0.5 mL 2 M lithium acetate (Sigma-Aldrich) and incubated on ice for 5 min. Cells were pelleted by centrifugation at 500 × *g*, 5 min, resuspended in 0.5 mL 0.4 M NaOH, and incubated on ice 5 min. For *E. invadens*, trophozoites or encysting parasites (3×10^5^) were pelleted by centrifugation at 500 × *g* for 5 min. Both yeast cells and *E. invadens* parasites were resuspended in NuPAGE LDS sample buffer (Life Technologies; Carlsbad, CA, USA). An additional step was required to lyse *E. invadens* cysts. Cysts (in NuPAGE LDS buffer) were also exposed to three cycles of freeze/thaw in liquid nitrogen.

All samples were heated for 5 min at 100°C and loaded onto a precast NuPAGE 12% Bis-Tris Gel (Life Technologies; Carlsbad, CA). The gels were electrophoresed at 180V for 60 min and proteins were transferred to PVDF membranes (Life Technologies) at 12V for 1.5 h in Towbin transfer buffer (25 mM Tris, 192 mM glycine, 20% (v/v) methanol). Prior to blocking, membranes were stained with Ponceau S reagent (Sigma-Aldrich) to record protein load.

The membranes were blocked with 5% (w/v) Blotting Grade powdered milk blocker (Bio-Rad Laboratories, Hercules, CA) and 0.5% (w/v) bovine gelatin (Sigma-Aldrich) in TBST (50 mM Tris, 150mM NaCl, 0.5% (v/v) Tween 20) for 35 min at 37°C. Membranes were incubated overnight at 4°C in primary antibodies (diluted 1:1000 in TBST). For yeast, the primary antibodies were horse radish peroxidase-conjugated α-poly-histidine tag antibody (Sigma Aldrich; St. Louis, MO; kind gift of Dr. Michael Sehorn, Clemson University), yeast-specific α-phosphorylated eIF2α antibody (Thermo-Fisher), or yeast-specific α-total eIF2α antibody (gift of Dr. Thomas Dever, NIH). For *E. invadens*, the primary antibodies were *Entamoeba*-specific α-phosphorylated eIF2α antibody [4] or α-total eIF2α antibody [4]. The membranes were washed in TBST for 45 min with 6 buffer changes. Membranes were incubated in commercially available horseradish peroxidase-conjugated goat anti-rabbit antibody (Thermo-Fisher, diluted 1:5000 in TBST) for 1 h at room temperature and extensively washed as described above. All blots were developed using a commercially available Enhanced ChemiLuminescence Western Blotting Detection system (ThermoScientific) according to the manufacturer’s instructions. Bands on the same membrane were quantified using scanning densitometry and Image J software (Version 1.51, NIH).

### Induction of Stage Conversion

To induce encystation, control and mutant trophozoites (6.5×10^6^) were pelleted by centrifugation at 500 × *g* for 5 min and resuspended in 47% (w/v) low glucose/serum free/high osmolarity encystation medium [4,7], supplemented with 50 mg/mL neomycin. Parasites were incubated at 25°C for either 48 h or 72 h and encystation was tracked by staining with Congo Red (Amresco; Solon, OH) and flow cytometry [19].

Excystation was induced as described [8]. Briefly, Trig Luc and EiIF2K-KD trophozoites were induced to encyst for 72 h. Parasites were then incubated in 13 mL ddH_2_O at 4°C overnight to lyse unencysted trophozoites. Cysts were enumerated using a Luna Automated Cell Counter (Logos Biosystems, Annandale, VA), pelleted by centrifugation at 500 × *g* for 5 min, and resuspended in 13 mL TYI-S-33 medium, 1 mg/mL bile salts (Sigma-Aldrich), and 40 mM NaHCO_3_, and incubated at 25°C for 2 h or 8 h. After incubation, cultures were iced for 8 min to detach any trophozoites from the glass culture tubes, pelleted by centrifugation at 500 × *g* for 5 min, resuspended in 1 mL of 1% (v/v) sarkosyl in PBS and incubated on ice for 30 min to lyse any trophozoites or immature cysts. The remaining detergent-resistant cysts were enumerated and the percent excystation was calculated by comparing total cysts remaining to the starting number of cells.

### Phagocytosis Assays

Phagocytosis assays were carried as previously described [20] with minor changes. Briefly, control or mutant trophozoites were rinsed once in PBS (GE Life Sciences) and twice in serum free TYI-S-33 medium (SFM). Trophozoites (2×10^5^) were resuspended in 150 μL SFM. Freshly isolated human red blood cells (hRBCS) were pelleted by centrifugation (2000 × *g* for 1 min) and rinsed once with PBS and twice with SFM and were resuspended at a concentration of 4×10^5^ cells/μL in SFM. hRBCs (2×10^7^) were added to the trophozoites and incubated at 25°C for 10 min. Samples were pelleted by centrifugation (2000 × *g* for 1 min), and undigested hRBCs were hypotonically lysed by washing twice with 1 mL of ice-cold ddH_2_O. Parasites were washed with 1mL ice-cold PBS, collected by centrifugation (2000 × *g* for 1 min) and lysed with 200 μL concentrated formic acid (Fisher). Phagocytosis was measured as the absorbance of heme in the lysate at 405 nm. Sample values were corrected using a formic acid blank.

### Adhesion Assays

Adhesion assays were carried as previously described [21] with minor changes. Briefly, control and mutant parasites were incubated with calcein-AM (Invitrogen) (5 μL/mL) for 30 min at 25°C. Chinese hamster ovary (CHO) cells (1.5×10^5^) were seeded into a 96-well plate and grown at 37°C for 24 h. CHO monolayers were fixed by incubating with 4% (v/v) paraformaldehyde in PBS for 10 min at 37°C. To inactivate paraformaldehyde, the CHO monolayer was incubated with 200 µL of 250 mM glycine for 15 min. Glycine was removed by rinsing with PBS. Calcein-AM labeled parasites were washed once with room temperature SFM and 5×10^4^ parasites were seeded onto the fixed monolayer of CHO cells. Parasites were incubated with fixed CHO cells in SFM for 30 min at 25°C. The media was carefully aspirated, and the cell layer was gently rinsed twice with room temperature PBS. The number adherent parasites was determined by measuring fluorescence at excitation and emission wavelengths of 495 and 525 nm, respectively, with a fluorimeter/plate reader (Model FLX800, BioTek Instruments, Winooski, VT).

### Induction of Oxidative stress

Control or mutant parasites were incubated with 4 mM or 1M H_2_O_2_ for 1 h at 25°C. Viability was assessed using trypan blue exclusion and quantified by an Automated Luna Cell Counter (Logos Biosystems).

### Statistical Analysis

All values are presented as means ± standard error of at least 3 separate trials. Means of treated groups were compared against the appropriate control and statistical analyses were performed using Graph Pad prism 9 (v9.0.0, San Diego, CA, US) with a one-way analysis of variance (ANOVA). *P* values of less than 0.05 were considered statistically significant. *P* values less than 0.01 or 0.001 were considered highly statistically significant.

### Ethics Statement

Whole blood was donated by a healthy adult volunteer, who provided oral consent, at Clemson University. The collection was approved by Clemson’s Institutional Biosafety Committee under safety protocol #IBC2018-12.

## Acknowledgements

The authors thank Dr. William Marcotte (Dept. of Genetics and Biochemistry, Clemson University) for the pYES-NT/C plasmid. The authors thank Dr. Ronald Wek (Indiana University School of Medicine) for the pYES2 plasmids containing live or dead human PKR kinase domains. The authors thank Dr. Lukasz Kozubowski, Dr. Mufida Ammar, and Rodney Colon-Reyes (Dept. of Genetics and Biochemistry, Clemson University) for their kind assistance with culturing yeast strains. The authors thank Dr. Michael Sehorn (Dept. of Genetics and Biochemistry, Clemson University) for anti-polyhistidine antibody. The authors thank Dr. Thomas Dever (National Institute of Child Health and Development, National Institutes of Health) for custom anti-total yeast eIF2α antibodies and Dr. Upinder Singh (Stanford University) for the Trigger plasmid. The authors thank Dr. Rooksana Noorai (Clemson University Genomics and Bioinformatics Facility) for helpful discussions on bioinformatic analysis of eIF2α kinases.

